# Error rates, PCR recombination, and sampling depth in HIV-1 Whole Genome Deep Sequencing

**DOI:** 10.1101/077313

**Authors:** Fabio Zanini, Johanna Brodin, Jan Albert, Richard A. Neher

## Abstract

Deep sequencing is a powerful and cost-effective tool to characterize the genetic diversity and evolution of virus populations. While modern sequencing instruments readily cover viral genomes many thousand fold and very rare variants can in principle be detected, sequencing errors, amplification biases, and other artifacts can limit sensitivity and complicate data interpretation. Here, we describe several control experiments and error correction methods for whole-genome deep sequencing of viral genomes. We developed many of these in the course of a large scale whole genome deep sequencing study of HIV-1 populations. We measured the substitution and indel errors that arose during sequencing and PCR and quantified PCR-mediated recombination. We find that depending on the viral load in the samples, rare mutations down to 0.2% can be reproducibly detected. PCR recombination can be avoided by consistently working at low amplicon concentrations.

---

Next-generation sequencing has become a standard tool for detection and genetic characterization of viruses. In most applications, the aim is to obtain one consensus sequence of the virus population in a sample with high viral titers (Gire *et al.*, 2014; Quick *et al.*, 2016; Zhou *et al.*, 2014). Plasma samples from HIV positive individuals often contain relatively few virus genomes and PCR amplification prior to sequencing is necessary. Moreover, minor variants within the sample are important to understand the biology and evolution of the virus, hence a single consensus sequence is not enough; the goal is to obtain a faithful representation of viral diversity despite PCR amplification. To this end, the amplification protocol must satisfy several criteria: the majority of template RNA molecules should be represented without bias; it should minimize mis-incorporations; and it should avoid PCR-mediated recombination to preserve linkage information. These requirements are partially conflicting – for instance more processive polymerases might generate less recombinants but also more frequent misincorporations – and finding an optimal compromise is essential.

Most projects characterizing diversity and minor variants in viral populations via deep sequencing have focused on specific amplicons that are short enough to be fully covered by a single (paired-end) read, typically using 454, IonTorrent, or Illumina technology (Bunnik *et al.*, 2011; Eriksson *et al.*, 2008; Hedskog *et al.*, 2010; Jabara *et al.*, 2011; Rozera *et al.*, 2009; Tsibris *et al.*, 2009). Such amplicon strategies provide great depth and allow a simple analysis workflow since every read covers the same locus on the viral genome. To our knowledge, Bimber *et al.* (2010) first demonstrated the feasibility of whole genome deep sequencing of SIV/HIV. Several similar strategies using different sequencing technologies have been developed since (Gall *et al.*, 2012; di Giallonardo *et al.*, 2014; Henn *et al.*, 2012; Ode *et al.*, 2015; Zanini *et al.*, 2016). Whole genome deep sequencing provides a comprehensive picture of the diversity within viral populations, but requires substantially more data processing steps than the single short amplicon design.

Here, we review different strategies for whole genome HIV deep sequencing with an emphasis on control experiments and computational tests necessary to quantify the accuracy of minor variant frequencies and the loss of linkage during sample preparation. We developed several such control experiments and checks as part of a previous study in which we sequenced HIV-1 genomes from a large number of plasma samples with low RNA HIV-1 copy number (Zanini *et al.*, 2016). In the following, we outline the critical steps during Illumina Sequencing of PCR amplified virus in order to obtain reliable linkage information and an assessment of actual sequencing depth starting from few template molecules.

## Extraction, amplification, and sequencing strategy

### RNA extraction

If virus specific PCR primers at conserved genomic regions are used, as in the majority of whole genome deep sequencing studies to date, RNA extraction must focus on maximal template yield rather than on purity of the extracted material, because non-viral nucleic acids that do not match the PCR primers will not be amplified. We used the RNeasy Lipid Tissue Mini Kit (Qiagen Cat. No. 74804) to extract total RNA, with two separate extractions each from 200 *µ*l of plasma and each with a double 50 *µ*l elution. Other extraction methods were tried, but we found that this method combined high sensitivity with relative ease of use. A similar kit, the Viral RNA Mini kit (Qiagen), was recently found to be performing well by the BEEHIVE Consortium (Cornelissen *et al.*, 2016).

### Amplification and primer design

We evaluated both gene-specific primers and random hexamers for cDNA synthesis. In agreement with others we found that gene-specific primers are more sensitive than random hexamers (Acevedo *et al.*, 2014). The majority of studies to date used specific primers to amplify a moderate number 1.5-3kb fragments of the HIV-1 genome, either using two nested PCRs (di Giallonardo *et al.*, 2014; Ode *et al.*, 2015) or a single PCR (Bimber *et al.*, 2010; Gall *et al.*, 2012; Zanini *et al.*, 2016).

When designing specific primers for viral amplification, the first critical decision is on the number and length of amplicons. Longer targets amplify less efficiently which typically leads to a more biased representation of quasi-species diversity in the sample. With more and shorter PCR targets, the initial viral RNA must either be split among more reactions with fewer template molecules per reactions, or amplicons must be carefully calibrated to ensure even amplification in a multiplex reaction. Furthermore, it is difficult to find many sufficiently conserved regions to accommodate a large number of primer pairs. Gall *et al.* (2012) and (Bimber *et al.*, 2010) proposed a set of 4 primer pairs ranging from 2 to 3.5 kb in length. In our study, we used 6 amplicons ranging from 1.5 to 2.1 kb in length. We tried to multiplex PCR targets no. 1, 3 and 5 in one PCR tube and targets 2, 4, and 6 in a second PCR tube, but we experienced problems with PCR efficiency and did not pursue this strategy further.

An alternative approach to using two specific primers per amplicon has been demonstrated recently by Berg *et al.* (2016). These authors have adapted the SMART cDNA synthesis strategy, which is commonly used for transcriptomics, to viral sequencing of HIV (HIV-SMART). Briefly, a special template-switching oligonucleotide (TSO) is used as a generic forward primer; using gene specific reverse primers or random hexamers together with the TSO, long stretches of HIV RNA can be amplified. However, the amplification specificity can be low such that a large fraction of reads don’t map to HIV. As SMART protocols become more sensitive and robust (Picelli *et al.*, 2014), this approach might also be useful for quantifying rare quasi-species variation.

Designing good primers for amplifying HIV-1 populations is challenging because of the extensive genetic variability of the virus. We proceeded with a two-step strategy. First, with the help of the software PrimerDesign-M (Brodin *et al.*, 2013; Yoon and Leitner, 2015), we designed primers that target conserved regions of HIV-1 genome, avoid primer dimers and hairpins, and have a uniform annealing temperature that allows all PCR reactions to be run in the same thermocycler. Although many of the initial primers were effective for diverse HIV isolates, we encountered difficulties amplifying fragment F5 covering the *gp120* gene. We were eventually able to design primers for this region but, as apparent from from the sequencing reads, we often amplified fewer template RNA molecules for *gp120* compared to the other amplicons. This difficulty appeared to be consistent across several different primer pairs, which suggests that it may be due to RNA secondary structure as the rev response element (RRE) is known to form a very stable RNA hairpin (Heaphy *et al.*, 1990). Interestingly, we found that adding random hexamers to the gene-specific primer increased sensitivity for some of the HIV-1 targets; random hexamers might help cDNA synthesis by destabilizing RNA secondary structures.

Ode *et al.* (2015) and di Giallonardo *et al.* (2014) used nested PCR in their whole genome deep-sequencing projects, which increases sensitivity and specificity of PCR amplification (Albert and Fenyö, 1990). We tested nested PCR strategies but settled on single round PCR since nested PCR increases of amplification biases and PCR-induced recombination. We used nested PCR only for control experiments and template quantification.

As the BEEHIVE project (Cornelissen *et al.*, 2016), we used Superscript III One-Step RT-PCR with Platinum Taq High Fidelity for cDNA synthesis and PCR. We found this reverse transcriptase mix to be efficient, sufficiently accurate and easy to use. We also tried reverse transcription using ThermoScript™ RT-PCR System and Superscript^®^ III First Strand System for RT-PCR but eventually chose the one-step approach, among other reasons, because it requires less handson time and thereby maximizes throughput.

To avoid PCR amplification bias and PCR recombination, the number of PCR cycles should be kept to a minimum (di Giallonardo *et al.*, 2013; Mild *et al.*, 2011) and library preparation protocols that require small amounts of input DNA are preferable. The Illumina Nextera XT technology (using a Tn5 transposase preloaded with adapters) allows reliable preparation of sequencing libraries for from as little as 500 pg of purified cDNA in around 2.5 ul. We routinely used 30 cycles of single PCR to amplify HIV-1 from patient samples, with a yield of around 0.5-5 ng of purified cDNA. If the DNA concentration after PCR was too low, we concentrated the amplicon in a vacuum centrifuge. If only some amplicons yielded measurable quantities of DNA, we nevertheless pooled all amplicons for library preparation and in many cases obtained useful sequencing data for most amplicons.

### Library preparation and sequencing

The Tn5 transposase fragments the 2kb amplicons into inserts of around 300 bp in length, with a distribution between 100 and 700bp. Since short inserts amplify preferentially during the final stages of library preparation and attach more readily to the flow-cell, the sequencing output tends to be dominated by short inserts. However, to study linkage between variants and optimal use the long reads delivered by the MiSeq platform (up to 2x300bp), long inserts are preferable. To obtain sequencing reads with inserts up to 700bp, we modified the standard protocol in two ways: (i) we reduced the ratio of transposase to input DNA by a factor of two, and (ii) replaced the bead size selection with the Blue Pippin (Sage Science) gel-based size selection. The latter allowed us to remove all reads with inserts shorter than 350bp and fully utilize the read length of the MiSeq sequencer. After size selection by the BluePippin, we obtained an even insert size distribution between 400 and 700 bps.

Despite its shorter read length, we preferred the 2x250 bp v2 MiSeq kit over the 2x300bp v3 kit since the 2x250bp v2 kit consistently delivered a larger number of high quality reads.

## Bioinformatics pipeline

The analysis of high-throughput sequencing reads from diverse species such as HIV-1 presents unique challenges and different approaches have been developed by different groups. Our analysis pipeline is available at https://github.com/neherlab/hivwholeseq (see Fig. S1 for an outline); here we will briefly address our approach to turn raw sequencing output into SNV frequencies, haplotypes, and linkage maps and compare it to alternative strategies.

The pipeline has two critical steps. The first challenge is mapping and assembling the reads in the presence of large natural variation (up to 10% including insertions and deletions). Many short read mappers such as BWA (Li and Durbin, 2009) use a fixed number of mismatches from the reference as a criterion for alignment success, but such a single, fixed threshold is difficult to apply to HIV-1 because within-population and between-population diversity varies greatly along the genome (Li *et al.*, 2015; Zanini *et al.*, 2016). Similarly, assemblers such as SPAdes (Bankevich *et al.*, 2012) can be inaccurate because they are designed to reconstruct a single haplotype and run into problems when assembling reads from a diverse population. We approached this problem by using a probabilistic mapper, Stampy (Lunter and Goodson, 2011) and by writing a custom assembly script that combines rough alignments to a HIV-1 reference sequence with *de novo* local realignment of the reads into a single contig followed by iterative refinement. Ode *et al.* (2015) also employed an iterative strategy combined with *de novo* assembly using vicuna (Yang *et al.*, 2012). (Cornelissen *et al.*, 2016) used the “iterative virus assembler” IVA (Hunt *et al.*, 2015). All these methods follow a similar strategy but rely do different degree on a reference for guiding assembly. While vicuna and IVA begin by de novo assembly with iterative merging and refinement, our strategy uses lenient mapping to a reference to scaffold reads before going through iterative refinement.

While filtering non-HIV reads is straightforward, detecting and removing sample cross-contamination is more subtle. Too strict filtering may lead to underestimation of “real” diversity whereas too little filtering results in inflated diversity estimates. Illicit reads might come from a number of sources, including (i) contamination of pre-PCR areas by cell lines used for viral production, (ii) cross-sample contamination during cDNA synthesis, PCR, or library preparation, and (iii) cross-talk between barcodes during multiplexed sequencing. The latter case is rare but not negligible: we observed about one in 10,000 reads coming from a different sample. While contamination can minimized by clean laboratory practices, cleaning the sequencing reads is essential. To this end, we compared each read to all consensus sequences from all patients studied and manually inspected the histograms of hamming distances of reads to the sample consensus for each genomic region. This way, contamination from a different HIV population can be detected and removed. However, in order for this strategy to work, genetically similar samples (e.g. from the same patient) have to be prepared on different days and in different PCR plates.

## Error rates in PCR and sequencing

To distinguish natural genetic variation in HIV-1 populations from sequencing errors, error profiles need to be known (Eriksson *et al.*, 2008; Orton *et al.*, 2015; Rosen *et al.*, 2012). Errors are generated during the reverse transcription, during PCR amplification (including any PCR during the library preparation), and during sequencing.

To assess the relative contributions of these sources of errors, we performed a series of control sequencing experiments. In each sequencing run, we included a 1% PhiX plasmid spike-in to evaluate the general performance of the run and the accuracy of the per-base phred quality scores. We performed similar control experiments for the combined amplification and sequencing pipeline by amplifying and sequencing HIV-1 plasmid DNA (2000 plasmid copies per PCR run). Errors were quantified by the frequency of nucleotides differing from the consensus sequence of the plasmid.

Using the phiX spike-in, we found that the quality scores reported by the Illumina instrument are reliable indicators of sequencing quality as shown in Fig. 1. We restricted analysis to positions with a phred score ≥ 30. This threshold was chosen such that the sequencing error rate would be lower than the error rate of the PCR (see below). Not much data is lost due to this filter since a large fraction of bases in a typical MiSeq run have phred qualities between 30 and 35. We also trimmed beginning and end of each read such that it doesn’t contain low-quality regions of size 10 with 2 or more bases with phred quality 20. The latter criterion is arbitrary but not critical, as the phred quality for each base is kept in the pipeline later on.

**Figure 1.**
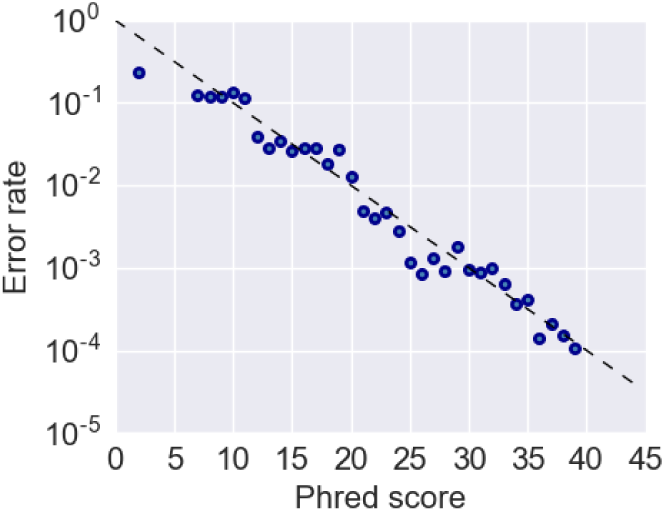
Quality scores reported by the MiSeq sequencing instrument are a reliable indicator of the per base error rate.

Ode *et al.* (2015) also observed that quality scores are useful for filtering sequencing errors, but used a slightly different strategy to remove errors. Instead of discarding base calls with qualities below a certain threshold, they averaged phred scores by nucleotide at each position. If the average score for a specific nucleotide was low (i.e. this nucleotide was predominantly found in low quality reads), this variant is classified as an error. While this strategy retains a larger number of bases, true rare variants might be lost among a large number of low quality reads. We routinely found that *>*70% of all bases have phred scores *>*30, such that masking all low quality bases was not a concern.

Fig. 2 characterizes sequencing and substitution errors. Panel A shows an error matrix for phiX control spike-in, i.e. how often nucleotide X is called when the correct nucleotide would have been Y – for instance, the first column, second row entry refers to the rate of mutation *C* → *A*. The error matrix shows a flat distribution across all mutations, which is expected given that we filter for high phred scores. In contrast, error matrices measured for the combined PCR amplification and sequencing protocol show a transition bias Fig. 2BC. Among transversions, *C* → *A* and *G → T* substitution errors are most frequent.

**Figure 2.**
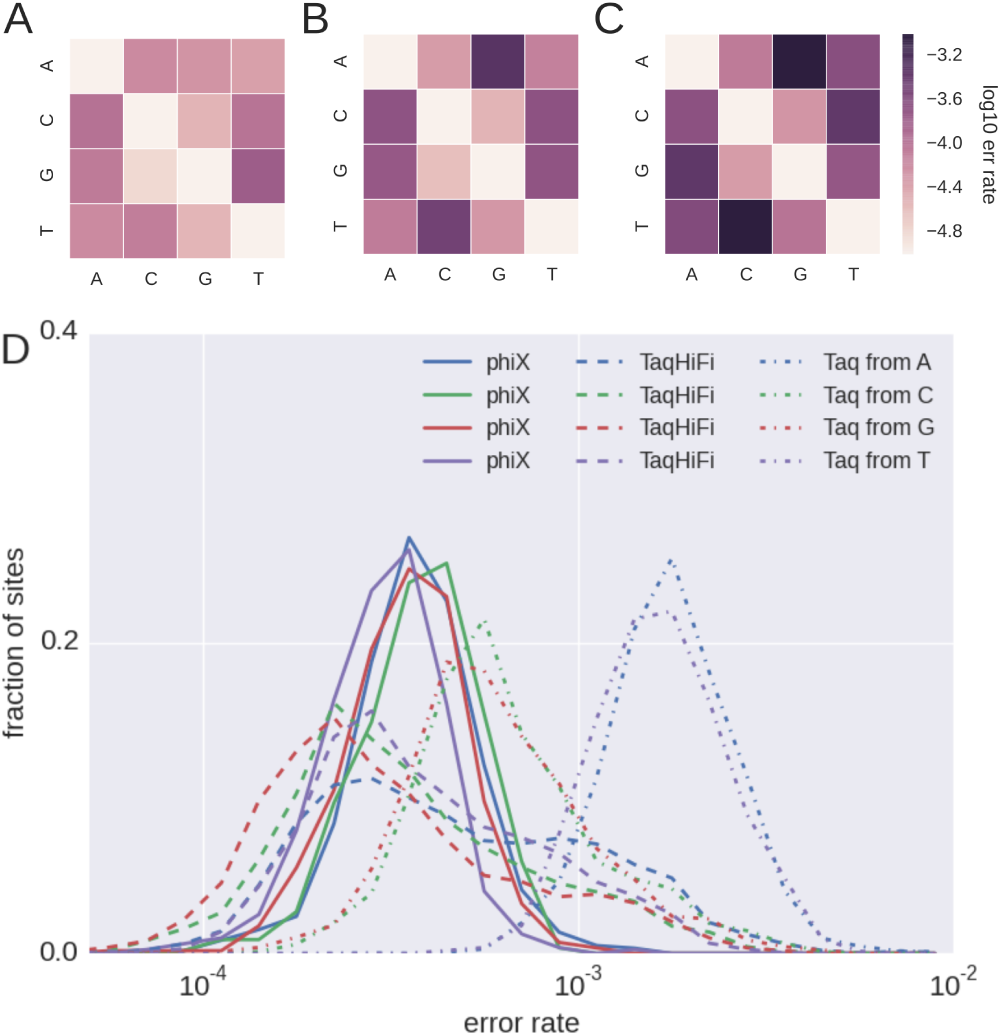
Substitution errors. To quantify substitution errors during PCR and sequencing, phiX DNA and virus plasmid NL4-3 were sequenced at high coverage. Panels A-C show the error matrices of the sequencing machine alone (A, phiX), sequencing and amplification errors using HiFi Taq polymerase (B) and regular Taq polymerase (C). Each row of the error matrices corresponds to a true template nucleotide. Entries in different columns indicate the rate at which the incorrect nucleotides are called. The sequencing errors alone (A) are evenly distributed and uniformly below 10^−3^. When the PCR step is included (B,C), most of errors are transitions. Panel D shows the distribution of the frequency of non phiX or NL4-3 variants at each site across the sequences. For TaqHiFi, error rarely exceed a frequency of 1/1000 while they are between 2-5 times higher for regular Taq.

Fig. 2D shows the distribution of error frequencies across sites with phred scores ≥ 30. The main peak in error frequencies for Platinum Taq Hifi agrees with that of phiX (~ a factor 2 lower, indicating run-to-run variation), but there is a pronounced tail at higher error rates presumably introduced during PCR. Nonetheless, the majority of sites have erroneous minor variants at frequencies about 3/10000. Exchanging the Platinum HiFi Taq polymerase for a normal Platinum Taq increased the error rate by a factor of 2-5.

In addition to filtering errors based on quality scores, some authors have employed clustering and haplotype reconstruction strategies to distinguish true variation from errors (di Giallonardo *et al.*, 2014; Rosen *et al.*, 2012; Zagordi *et al.*, 2010). Such approaches are most useful to study populations with limited recombination, e.g. HIV during early infection, influenza virus, or HCV. During chronic HIV infection, however, recombination results in a rapid decay of linkage along the genome (Neher and Leitner, 2010; Zanini *et al.*, 2016) and haplotype reconstruction might underestimate the genetic diversity of the population.

In summary, the majority of errors arise during PCR at a rate of about 0.1% after filtering low quality base calls, as has also been reported in other studies (di Giallonardo *et al.*, 2013; Mild *et al.*, 2011).

## Linkage

Previous studies have shown that *in vitro* recombination depends strongly on the concentration of the amplicons (di Giallonardo *et al.*, 2013; Mild *et al.*, 2011) as well as on the RT-PCR conditions (Fang *et al.*, 1998; Shao *et al.*, 2013). We therefore optimized the sequencing library preparation to work with as little input DNA as possible (typically one nanogram). This allowed us to amplify all samples with a single PCR (as opposed to nested PCR), which strongly reduced PCR-induced recombination (see below). If nested PCR is necessary to obtain enough material for sequencing, however, several improvements over manufacturers specifications have been shown to reduce *in vitro* recombination from 10% or more recombinant reads to fewer than 1%: longer extension times during cDNA synthesis, a higher concentration of primers, careful quantification of input DNA into the second round of PCR, and potentially skipping the final extension after PCR (Fang *et al.*, 1998; di Giallonardo *et al.*, 2013; Shao *et al.*, 2013).

We tested both nested and single PCR for our study on a 50/50 mix of two HIV-1 laboratory strains in order to quantify the rate of PCR-induced recombination. We found a significant fraction of recombinant reads after nested PCR (around 10%) but almost no recombinant reads after single PCR. The loss of linkage due to recombination can be quantified by the linkage disequilibrium measure *D′*. For two variants at frequency *p*_1_ and *p*_2_, *D′* is defined as

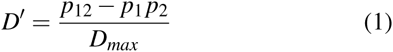

where *p*_12_ is the fraction of reads that carry both mutations. The denominator *D*_*max*_ normalizes for differences in *p*_1_ and *p*_2_ and is given by min(*p*1*p*2, (1 – *p*_1_)(1 – *p*_2_) if *D′* < 0 and min(*p*1(1 – *p*2), (1 – *p*_1_)*p*_2_ if *D′* > 0 (Hartl and Clark, 2007). Linkage disequilibrium after nested PCR (PCR2) decreases within 200bp, a telltale sign of in vitro recombination (see Fig. 3), but the curve is flat for single PCR (PCR1) indicating that linkage is not lost during the PCR1.

**Figure 3.**
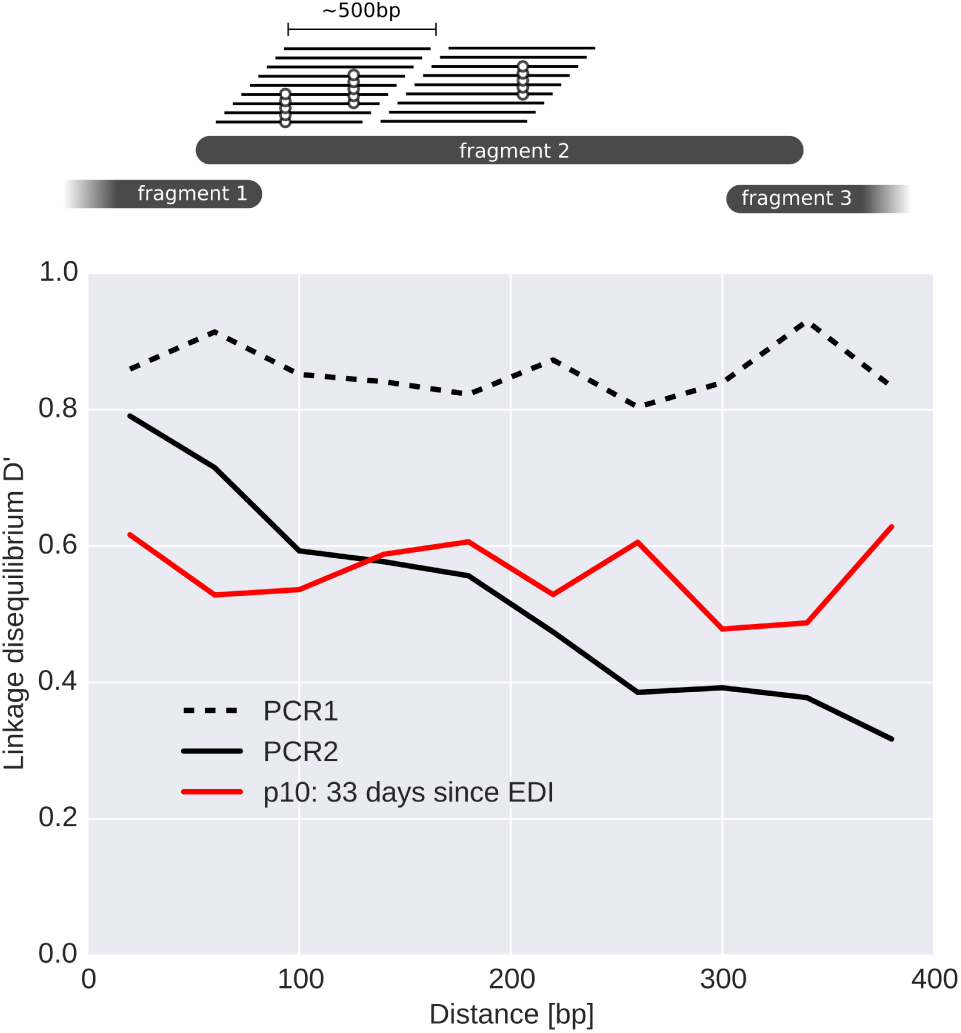
In vitro recombination. Short read sequencing data can be used to quantify linkage between two mutations if the corresponding positions are frequently sequenced on the same read. In our project, we can measure linkage within fragments up to a distance of about 500 bps, as illustrated in the top panel. The lower panel quantifies the effect of artefactual PCR recombination that potentially compromises linkage information. The lack of decay of linkage disequilibrium *D′* with distance in the PCR1 control and the early patient sample (33 days since the estimated date of infection (EDI)) demonstrates that one round of PCR at low template concentrations does not result in substantial in vitro recombination. A second nested PCR at higher template concentration results in considerable PCR recombination (PCR2).

Linkage controls are usually artificial mixtures of plasmids or almost homogeneous virus cultures. Some patient samples, however, provide an almost equally well-defined control.

Among the samples investigated in (Zanini *et al.*, 2016), one sample was from an individual who was apparently infected with more than one virion from the same donor very shortly prior to the collection of the first sample. The sample showed clearly two peaks of SNV frequencies suggesting that three variants dominated the initial population. Linkage between those variants was almost complete as shown by the lack of LD decay in Fig. 3. This is an internal confirmation that our single-PCR amplification strategy was not affected by in-vitro recombination.

## Indels

While Illumina’s sequencing technology generates much fewer indel errors than most other NGS technologies, such errors still occur during reverse transcription, PCR and sequencing. When sequencing libraries produced from homogeneous plasmids, we observe deletions and insertions at frequencies of about 1 in 105 sequenced bases (see Fig. 4 and Fig. 5). Larger deletion rates are observed at homopolymeric tracts, suggesting that slippage during PCR is a major source of these errors (see Fig. 5). Libraries produced from largely homogeneous viral cultures have a ten fold higher frequency of insertions and deletions than plasmid samples, possibly indicating a higher rate of indels during the RT step. Furthermore, the insertions of length 2 and three are much less common relative to those of length 1.

**Figure 4.**
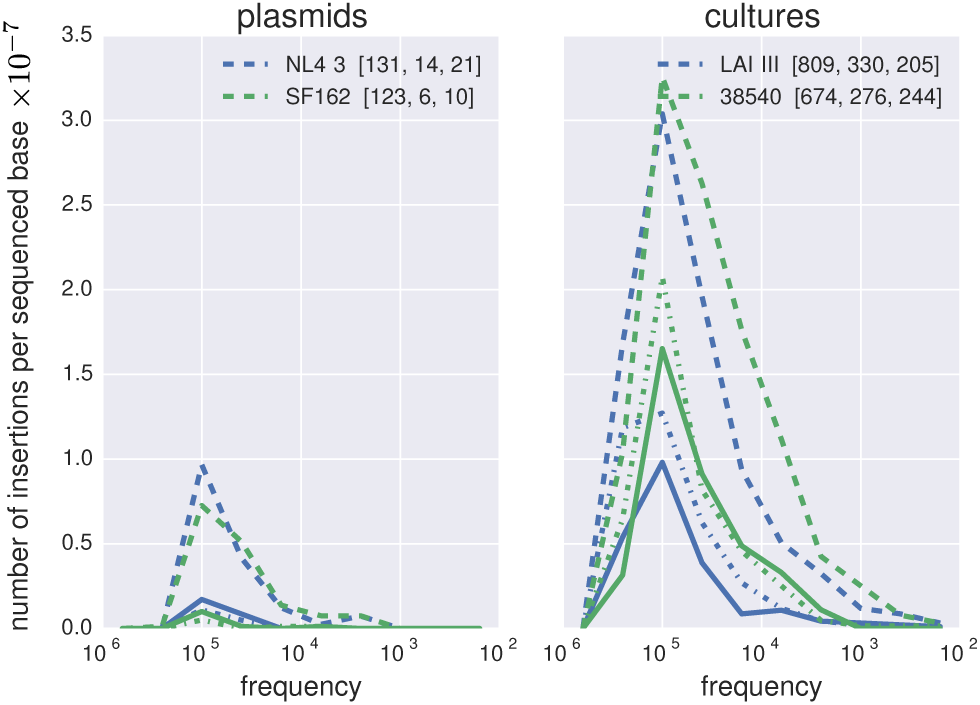
Insertion errors. Panel A shows the number of insertion observed at different frequencies relative to reference plasmids, normalized to the total number of bases sequenced. Dashed lines corresponds to insertions of length 1, dash-dotted to insertions of length 2 and solid to length 3. Total number of observed insertions in each experiment are given in the legend (length 1,2,3). Panel B shows the equivalent results for virus cultures. Insertions of length one are about three times more common in cultured virus than in plasmids. Two base or longer insertions were very rare in sequencing reads derived from plasmids, but were observed at appreciable frequencies in virus cultures.

**Figure 5.**
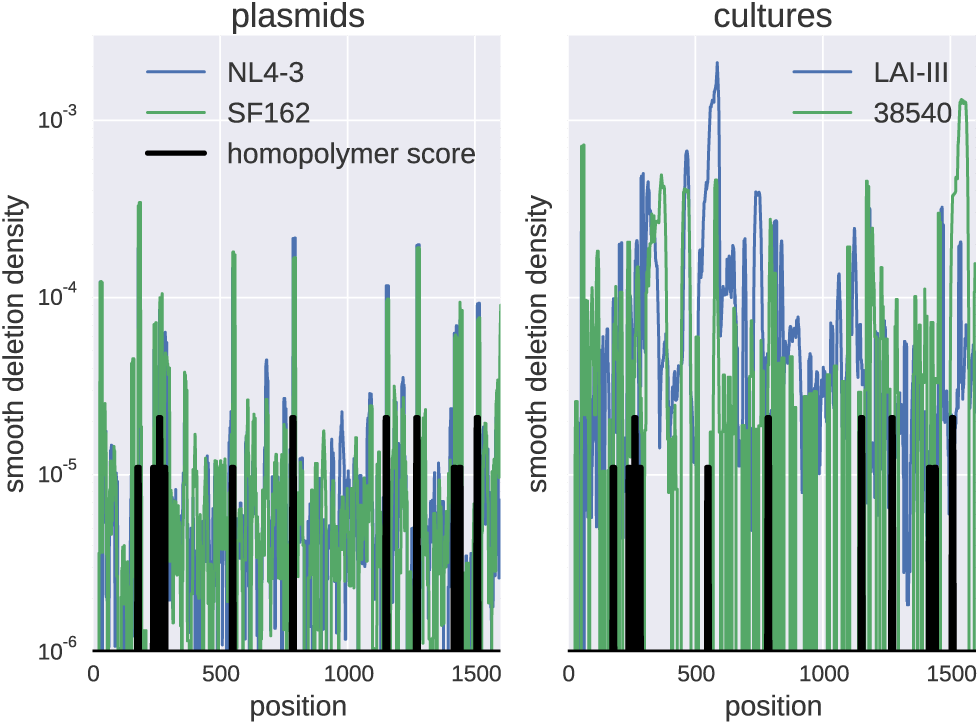
Deletion Errors. Panels A and B show the density of deletions along fragment 2 smooth with a 10 base window for plasmid controls and virus cultures, respectively. Deletions are more evenly distributed in virus cultures but still correlate with homopolymer densities. When sequencing plasmids, deletion errors localize strongly at homopolymer tracts.

## Depth and the accuracy of variant frequency estimates

Several studies have quantified the accuracy at which variant frequencies can be measured with deep sequencing. The predominant strategy is mixing of readily distinguishable viral genomes in defined proportions. Ode *et al.* (2015) have found that minor variants at 1% can be reliably detected. Seifert *et al.* (2016) used a five virus mix to show that frequencies of different viruses are reproducible to within one percentage point.

These tests used several thousand molecules as input for RT-PCR. In low viral load clinical samples, the accuracy of variant frequencies is primarily determined by the number of available template molecules (Iyer *et al.*, 2015). It is therefore important to quantify template input, PCR efficiency and bias for each sample.

In our whole genome deep-sequencing project, we used fragment 4 to estimate the template input by limiting dilution since fragment 4 amplified most reliably. 10% of the sample for fragment 4 was used for duplicate 10-fold dilution series by nested PCR. The series of positive and negative PCR reaction was used to estimate the number of templates based on a Poisson model of template sampling. The estimates of template input correlated well with viral load measurements made at the time of sampling (rank correlation ρ = 0.7). The median efficiency (the ratio between estimated template input and viral load) was 30% (Zanini *et al.*, 2016).

In addition to the low and variable template input, RT-PCR efficiency and biases can vary from fragment to fragment. Hence a global estimate of the of template numbers is sometimes not sufficient: ideally SNV frequency accuracy should be determined for every PCR reaction. This is difficult since determining those frequencies is the very goal of the sequencing experiment. However, overlapping PCR fragments allow for a built-in control: Every SNV in the overlap region is amplified and sequenced twice in completely independent reactions illustrated in Fig. 6. Concordance between these independent estimates can be used to estimate the effective template number contributing to each fragment.

**Figure 6.**
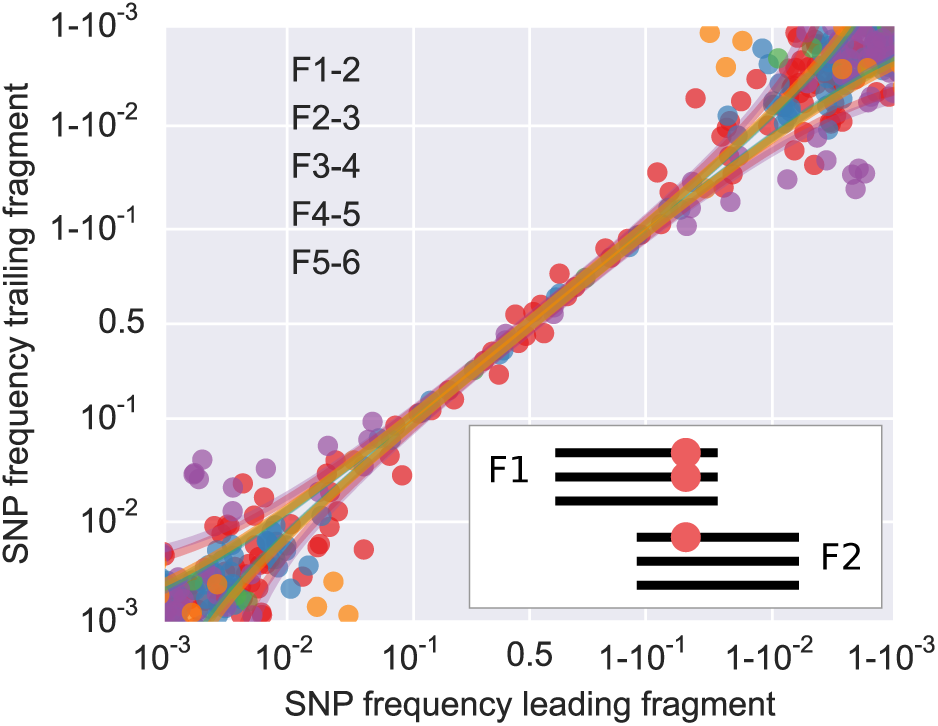
Estimating depth from SNV frequency overlap. The strategy to amplify overlapping fragments allows estimation of template numbers and the accuracy of SNV frequencies by comparing SNV frequencies in fragment overlaps. The deviation between frequencies estimated in trailing and leading fragment can be used to obtain an effective number of templates contributing to sequencing reads for each fragment – provided there is sufficient diversity in the overlap. Reproduced from (Zanini *et al.*, 2016).

Considering one overlap first, we can model the deviations of SNV frequency estimates as follows. Let the true frequency of the variant be *x* and assume that the leading and trailing fragment are presented in the sequencing library by *t*_1_ and *t*_2_ effective templates (we will discuss the effective nature of these numbers below). If the templates are sampled independently of the variant in question, the number of templates carrying the variant will be binomially distributed with means *xt*_1_ and *xt*_2_ and variances *x*(1 – *x*)*t*_1_ and *x*(1 – *x*)*t*_2_ in the leading and trailing fragment, respectively. If we further assume that many templates of either variant are captured, the binomial distribution can be approximated by a Gaussian distribution. The difference between the estimated frequencies in the leading and trailing fragment 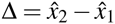 is then given by

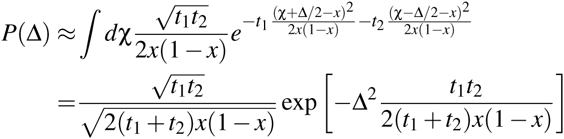

where the integral is over the average frequency of the variant in both segments. Hence the variance of the SNV frequency discordance ∆ is given by σ^2^ = (*t*_1_ +*t*_2_)*x*(1 – *x*)*/t*_1_*/t*_2_ which is dominated by the smaller of *t*_1_ and *t*_2_. The ratio ∆*/*(*x*(1 – *x*)) is expected to have a variance on the order of (*t*_1_ + *t*_2_)*/t*_1_*/t*_2_ and we estimate the template numbers by averaging ∆*/*(*x*(1 – *x*)) over different SNV *i* in the overlap. Approximating *x* by the average estimate 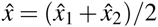, we obtain

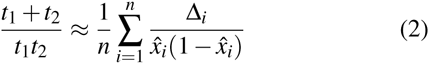

where the sum runs over different SNVs in the overlap. Given enough diversity, we can estimate this variance for each of the fragment overlaps. To stabilize estimates that are driven by few data points, we add several pseudo-SNVs (three) each of which contributes the inverse of the template number estimated from limiting dilutions.

### Approximate solution of the under-determined problem

For *k* fragments we only have *k* – 1 overlaps and solving for the *k* template numbers is therefore an ill-posed problem. However, a simple heuristic lower bound to template numbers works well. Since the variance observed in one overlap is dominated by the smaller of the two template numbers, we can start with the overlap with the smallest variance and solve for the leading and trailing template numbers assuming they are equal. From there, we work our way outward to noisier overlaps, always assigning the leftover variance to the outer fragment. Alternatively, one could solve for the template numbers along with a regularization that keeps template numbers as similar as possible.

### Limitations

For the method to work, we require several SNVs in the overlap region. The method is therefore unsuitable for short overlaps and homogeneous samples (e.g. very early samples) and even in cases with several SNVs in the overlap, the resulting estimates are noisy. Nevertheless, fragment specific indicators of input and amplification success are valuable controls; visual inspection of SNV frequency concordance is helpful to spot poorly performing fragments even if a quantitative analysis via equation (2) is not feasible.

While overlapping amplicons have value as intrinsic control, overlaps should not be excessively long. Once they become longer than the length of the sequencing inserts, reads can no longer be confidently assigned to one or the other fragment unless reads begin or end at a primer site or are barcoded in a fragment specific manner.

### Discussion

With high throughput sequencing technologies millions of viral genomes can be sequenced within hours or days. However, harnessing this sequencing capacity and obtaining accurate and interpretable results remains a challenge as errors and biases need to be carefully controlled. Sequencing technologies are changing rapidly and each new technology has different error profile and input DNA requirements. The Illumina MiSeq platform has comparatively long reads, low error rates, low DNA input requirements, and fast turn around time; it is therefore well suited for virus sequencing projects.

The main limitation when sequencing low viral load HIV-1 plasma samples is the limited number of HIV-1 genomes and efficient amplification of long segments. Similar challenges have been encountered efforts in recent efforts to sequence Zika virus genomes from clinical samples (Zibraproject, 2016). Quantification of template input is crucial for the interpretation of the results, in particular when intra-host variation is of interest.

We used limiting dilution together with viral load measurements and overlapping amplicons to assess the number of input templates (Zanini *et al.*, 2016). Another method to quantify template input and PCR amplification biases is the use of primerIDs – also called unique molecular identifiers – where a unique random sequence tag is introduced in the reverse transcription reaction to label the starting molecule. If this tag is subsequently amplified and sequenced in multiple reads, biases and sequencing errors can be corrected by only considering consensus sequences sharing one tag (Jabara *et al.*, 2011). However, primerID methods are not applicable when the amplicons are too long to be sequenced in one read, since shearing or tagmentation decouples the sequence from the ID. Digestion-circularization protocols have been recently put forth as a way to overcome the latter issue but typically require large amounts of input nucleic acids (Acevedo *et al.*, 2014; Hong *et al.*, 2014). Furthermore, primerID methods might introduce additional amplification biases and not necessarily improve the accuracy of variant frequency estimates (Brodin *et al.*, 2015; Seifert *et al.*, 2016).

Long-read sequencing platforms such as the RS-II (Pacific Biosciences) and the MinION (Oxford Nanopore) might soon make fragmentation of 2-4kb amplicons unnecessary. The current library preparation protocols for these long read technologies require large amounts of DNA input, making them unsuitable for low viral load samples. For instance, in order to adapt our protocol to long-read sequencing platform, we would need to add a second (nested) PCR that generates high rates of in vitro recombination (see above). Another important concern is that the error rates of single molecule sequencing techniques are high, such that error correction methods need to be applied. Despite these hurdles, circular consensus sequences from Pacific Biosciences have been recently used to characterize whole env diversity in high viral load samples (Laird Smith *et al.*, 2016). With increasing read length, reduced error rates, and innovative error correction schemes, whole genome viral sequencing in one contiguous read might soon become possible.

## Acknowledgements

We are grateful to Lina Thebo and Christa Lanz for excellent technical support. This work was supported by the European Research Council through grant Stg. 260686 and the Swedish Research Council trough grant K2014-57X-09935.

## Appendix: Supplementary material

**Figure S1.**
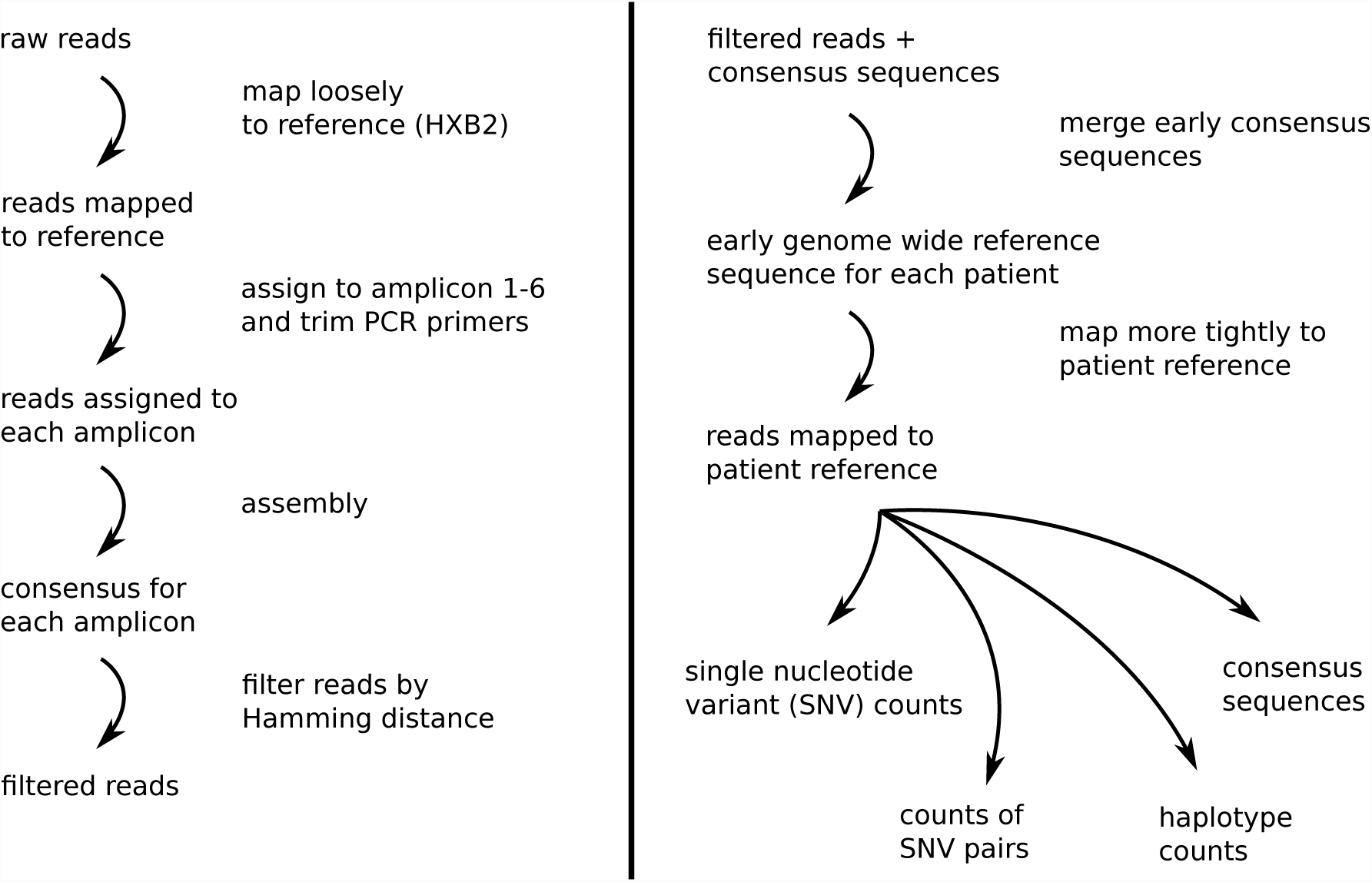
Outline of the data analysis workflow for HIV whole genome deep sequencing as performed in our previous study (Zanini *et al.*, 2016) (adapted from https://github.com/neherlab/hivwholeseq).

